# PHGDH supports liver ceramide synthesis and sustains lipid homeostasis

**DOI:** 10.1101/838482

**Authors:** Yun Pyo Kang, Aimee Falzone, Min Liu, James J. Saller, Florian A. Karreth, Gina M. DeNicola

**Affiliations:** Department of Cancer Physiology, H. Lee Moffitt Cancer Center and Research Institute, Tampa, FL, USA; Proteomics and Metabolomics Core Facility, H. Lee Moffitt Cancer Center and Research Institute, Tampa, FL, USA; Department of Anatomical Pathology, H. Lee Moffitt Cancer Center and Research Institute, Tampa, FL, USA; Department of Molecular Oncology, H. Lee Moffitt Cancer Center and Research Institute, Tampa, FL, USA

**Keywords:** PHGDH, serine, ceramide, triacylglycerol, mouse model

## Abstract

**Background:** D-3-phosphoglycerate dehydrogenase (PHGDH), which encodes the first enzyme in serine biosynthesis, is overexpressed in human cancers and has been proposed as a drug target. However, whether PHGDH is critical for the proliferation or homeostasis of tissues following the postnatal period is unknown.

**Methods:** To study PHGDH inhibition in adult animals, we developed a knock-in mouse model harboring a PHGDH shRNA under the control of a doxycycline-inducible promoter. With this model, PHGDH depletion can be globally induced in adult animals, while sparing the brain due to poor doxycycline delivery.

**Results:** We found that PHGDH depletion is well tolerated and no overt phenotypes were observed in multiple highly proliferative cell compartments. Further, despite detectable knockdown, liver and pancreatic function were normal. Interestingly, diminished PHGDH expression in the liver reduced serine and ceramide levels without increasing the levels of deoxysphingolipids. Further, liver triacylglycerol profiles were altered, with an accumulation of longer chain, polyunsaturated tails upon PHGDH knockdown.

**Conclusions:** These results suggest that dietary serine is adequate to support the function of healthy, adult murine tissues, but PHGDH-derived serine supports liver ceramide synthesis and sustains general lipid homeostasis.

## Background

The amino acid L-serine is derived from the diet, protein degradation, hydroxymethylation of L-glycine, and/or *de novo* biosynthesis. In the first step of de novo L-serine biosynthesis, D-3-phosphoglycerate dehydrogenase (PHGDH) catalyzes the formation of 3-phosphohydroxypyruvate from the glycolytic intermediate 3-phosphoglycerate. PHGDH activity is increased in many cancer cells as a consequence of genomic amplification, transcriptional upregulation, posttranslational modification, and allosteric regulation (DeNicola et al., 2015; Ding et al., 2013; Hitosugi et al., 2012; Locasale et al., 2011; Ma et al., 2013; Ou et al., 2015; Possemato et al., 2011; Tameire et al., 2019). Cancer cells with increased PHGDH activity are more dependent on PHGDH for proliferation, suggesting PHGDH is an attractive target for cancer therapy. Indeed, multiple PHGDH inhibitors have been developed, although none have reached the clinic to date.

While the contribution of PHGDH and serine to cancer cell metabolism has been well studied (Diehl et al., 2019; Labuschagne et al., 2014; Maddocks et al., 2013; Maddocks et al., 2016; Maddocks et al., 2017; Pacold et al., 2016; Sullivan et al., 2019; Ye et al., 2014; Ye et al., 2012), less is known about the importance of PHGDH in normal tissues. Whole animal PHGDH deletion is embryonic lethal in mice as a consequence of overall developmental retardation and brain defects (Yoshida et al., 2004). Brain-specific deletion dramatically reduces L-serine and D-serine levels in the cerebral cortex and hippocampus (Yang et al., 2010), leads to the development of post-natal microcephaly (Yang et al., 2010), and results in the accumulation of toxic deoxysphingolipids in the hippocampus (Esaki et al., 2015). Targeted deletion of PHGDH in endothelial cells is similarly lethal shortly after birth as a consequence of vascular defects due to compromised heme synthesis and mitochondrial function (Vandekeere et al., 2018). By contrast, PHGDH deletion in adipocytes presents with no overt phenotype, but improves glucose tolerance upon diet-induced obesity (Okabe et al., 2018). However, whether PHGDH is critical for the proliferation or homeostasis of other tissues following the postnatal period is unknown.

To study how PHGDH inhibition affects the functions of normal tissues in adult animals, we developed a knock-in mouse model harboring a PHGDH shRNA under the control of a doxycycline-inducible promoter. With this model, PHGDH depletion can be globally induced in adult animals, while sparing the brain due to poor doxycycline penetration. We find that PHGDH depletion is well tolerated in multiple highly proliferative cell compartments, with no overt phenotypes observed following knockdown. Further, PHGDH knockdown leads to a reduction in ceramide levels and an increase in triglyceride long chain (LC) poly unsaturated fatty acid (PUFA) content. These results suggest that dietary serine is adequate to support the function of healthy, adult murine tissues.

## Methods

### Reagents

HPLC grade Chloroform (Sigma Aldrich, 650498-1L) and Methanol (Sigma Aldrich, 34860-1L-R) were obtained from Sigma Aldrich. HPLC grade water (W5-1) was from Fisher Scientific. The following internal standards were used in targeted lipidomics analyses: Cer/Sph Mixture II (contains Cer(d18:1/12:0), cat #LM6005, Avanti Polar Lipids), D_7_-Sphinganine (860658, Avanti Polar Lipids), D_3_-Deoxysphinganine (860474, Avanti Polar Lipids), C12-doxCer [Cer(m18:1/12:0)] (860455P-1mg, Avanti Polar Lipids), and C12-dihydro-doxCer [Cer(m18:0/12:0)] (860481P-1mg, Avanti Polar Lipids). The following internal standards were used in metabolomics analyses: D_3_-Serine (DLM-582-0.1, Cambridge Isotope Laboratories), and the METABOLOMICS AMINO ACID MIX STANDARD (contains ^13^C_3_, ^15^N-Serine, cat #MSK-A2-1.2, Cambridge Isotope Laboratories)

### shRNA validation

NIH3T3s (ATCC) and Lenti-X 293T cells (Clontech) were grown in DMEM supplemented with 10% FBS. shRNAs were cloned into the LT3GEPIR vector (Fellmann et al., 2013) for shRNA testing. Lentivirus was produced in 293T cells using helper plasmids pCMV-dR8.2 dvpr (addgene # 8455) and pCMV-VSV-G (addgene #8454). NIH3T3 cells were infected with lentivirus encoding doxycycline-inducible shRNA constructs at an MOI of 0.2 for single-copy integration. Cells were treated with 1μg/mL doxycycline for 6 days and PHGDH expression determined by western blot. shRNA #3 (targeting 5’-ACCTGAACTAATACCTAGTAA-3’) was selected and cloned into the col1A1 targeting vector cTGME for ESC targeting.

### Mice

To generate shPHGDH mice, C10 murine ES cells (Beard et al., 2006) were targeted by recombination-mediated cassette exchange as previously described (Premsrirut et al., 2011) and selected with hygromycin. Positive clones were screened by PCR and injected into blastocysts. Mice expressing a Renilla Luciferase control shRNA (shREN) (Premsrirut et al., 2011) and ROSA26-CAGs-rtTA3 mice (Dow et al., 2014) were obtained from Dr. Lukas Dow (Weill Cornell Medicine). Mice were housed and bred in accordance with the ethical regulations and approval of the IACUC (protocol #IS00003893R). Mice were maintained on a mixed C57B6/129 background. shPHGDH; R26-CAGs-rtTA3 and shREN; R26-CAGs-rtTA3 mice were given a doxycycline-containing diet (200 ppm, Envigo), which was replaced weekly.

### Blood counts

Blood was collected from submandibular vein into Eppendorf tubes containing EDTA. Samples were completely mixed and stored at room temperature until analysis. All blood samples were analyzed within 4 hours of collection. Complete blood counts were analyzed with the Procyte Dx Hematology Analyzer (IDEXX).

### Oral glucose tolerance test

Mice were placed on doxycycline diet for approximately 100 days before use. Mice were fasted overnight, followed by oral administration of 2g/kg D-glucose solution. Blood was serially sampled at the indicated time points and glucose levels determined with the OneTouch Ultra Mini Blood Glucose Monitoring System.

### Immunohistochemistry

Tissues were fixed in 10% formalin overnight before embedding in paraffin and sectioning by IDEXX BioAnalytics. Sections were deparaffinized in xylene, followed by rehydration in a graded alcohol series. Heat-mediated antigen retrieval (microwave, 12.5 minutes on high) was performed in 10 mM citrate buffer (pH 6.0). Endogenous peroxidase activity was quenched with 3% hydrogen peroxide in tap water for 5 minutes. Immunohistochemical staining was performed with the ImmPRESS HRP anti-rabbit kit according to manufacturer’s instructions (Vector Labs), followed by incubation with DAB substrate (Vector Labs). The following antibodies were used: Ki-67 (1:400; Cell Signaling, 12202).

### LC-MS based Lipidomics

The liver tissue samples were homogenized with pre-chilled BioPulverizer (59012MS, BioSpec) and then placed on dry ice. The chloroform:methanol extraction solvent (v:v = 1:2) containing 5 nM Cer(m18:1/12:0), 5 nM Cer(m18:0/12:0), 12.5 nM D_3_-Deoxysphinganine, and 50 nM Cer(d18:0/12:0) internal standards was added to homogenates to meet 50 mg/mL. The samples were then sonicated in ice cold water using Bioruptor^TM^ UCD-200 sonicator for 5 min (30s sonication and 30s rest cycle; high voltage mode). For serum samples, 75 μL of chloroform:methanol extraction solvent (v:v = 1:2) containing 5 nM Cer(m18:1/12:0), 50 nM of Cer(d18:0/12:0), and 12.5nM D_3_-Deoxysphinganine internal standards were added to 20 μL of mouse serum. After shaking (1400 rpm, 20°C, 5 min), the extracts were cleared by centrifugation (17,000g, 20°C, 10 min), and the lipids in the supernatant were analyzed by LC-MS.

The HPLC condition was adapted from a previous study (Kang et al., 2014). Chromatographic separation was conducted on a Brownlee SPP C18 column (2.1 mm x 75mm, 2.7 µm particle size, Perkin Elmer, Waltham, MA) using mobile phase A (100% H2O containing 0.1% formic acid and 1% of 1M NH4OAc) and B (1:1 acetonitrile:isopropanol containing 0.1% formic acid and 1% of 1M NH4OAc). The gradient was programmed as follows: 0-2 min 35% B, 2-8 min from 35% B to 80% B, 8-22 min from 80% B to 99% B, 22-36 min 99% B, 36.1-40 min from 99% to 35% B. The flow rate was 0.400 mL/min. The parallel reaction monitoring (PRM) approach was applied for quantification of deoxysphingolipids in positive ESI mode. The MS and MS/MS m/z values of deoxysphingolipids for PRM analysis were adapted from a previous study (Esaki et al., 2015) (**Supplementary Table 1**). The collision energies of deoxysphingolipid species for PRM approach were set as following: 40 for doxCer and dihydro-doxCer; 15 for deoxysphinganine. For non-targeted lipidomics, the data dependent MS^2^ scan conditions were applied: The scan range was from m/z 70-1000, resolution was 120, 000 for MS and 30,000 for DDMS^2^ (top 10), AGC target was 3E6 for full MS and 1E5 for DDMS^2^, allowing ions to accumulate for up to 200 ms for MS and 50 ms for MS/MS. For MS/MS, the following settings are used: isolation window width 1.2 m/z with an offset of 0.5 m/z; stepped NCE at 10, 15 and 25 a.u., minimum AGC 5E2, and dynamic exclusion of previously sampled peaks for 8 s.

For targeted lipidomics, deoxysphingolipid and ceramide peaks identified by MS^2^ were manually integrated using Thermo Xcaliber Qual Browser. The quantification was based on previous methods (Esaki et al., 2015; Schutzhold et al., 2016). For non-targeted lipidomics, lipid peaks including triacylglycerols and ceramides were identified, aligned, and exported using MS-DIAL (Tsugawa et al., 2015). The data were further normalized to the median value of total lipid signals. Only lipids fully identified by MS^2^ spectra were included in the analysis.

### Serum serine quantification by GC-MS

50 μL of serum was extracted at −80°C for 15 min with 450 μL of 88.8% MeOH containing 205 μM D_3_-Serine. Following centrifugation (17,000g, 4°C, 20 min), 100 μL of supernatant was transferred to a new Eppendorf tube, and then dried by centrifugation under vacuum (SpeedVac, Thermo Scientific). The dried pellets were further derivatized as previously described (Carey et al., 2015). Briefly, the pellets were derivatized by 50 μL of methoxylamine hydrochloride (40 mg/mL in Pyridine) at 30°C for 90 min. The derivatized solution was then mixed with 70 μL of Ethyl-Acetate in a glass vial, the mixture was further derivatized with 80 μL of MSTFA + 1%TMCS solution at 37°C for 30 min. The final derivatized solution was then analyzed by GC-MS as previously described (Kang et al., 2019) using a MS scan range from 50 to 600 m/z. The derivatized serine (3TMS, 306 m/z) and D_3_-Serine (3TMS, 309 m/z) peaks were extracted and integrated manually using Agilent MassHunter Qualitative Analysis Software (Version B.07.00). The quantification was based on previous methods (Bennett et al., 2008).

### Liver serine quantification by LC-MS

After pulverization of liver tissue by pre-chilled BioPulverizer (59012MS, BioSpec), the extraction solvent (80% MeOH containing 2.49 μM ^13^C_3_, ^15^N-Serine, −80C) was added for a final concentration of 50 mg tissue/mL and incubated for 24 hr at −80°C. The metabolite extracts were centrifuged (17,000g, 4°C, 20 min), and the supernatant was analyzed by LC-MS as previously described (Kang et al., 2019). Data was acquired in ESI-positive mode. For targeted quantification of serine, ^12^C-Serine and ^13^C_3_, ^15^N-Serine peak areas were manually integrated by EL-Maven (Version 0.6.1). Quantification was based on previous methods (Bennett et al., 2008).

### Immunoblotting

Tissue lysates were prepared by dounce homogenization in RIPA buffer (20 mM Tris-HCl [pH 7.5], 150 mM NaCl, 1 mM EDTA, 1 mM EGTA, 1% NP-40, 1% sodium deoxycholate) containing protease inhibitors (Roche complete). Protein concentrations were determined by the DC protein assay (Bio-Rad). Lysates were mixed with 6X sample buffer containing β-ME and separated by SDS-PAGE using NuPAGE 4-12% Bis-Tris gels (Invitrogen), followed by transfer to 0.45μm Nitrocellulose membranes (GE Healthcare). The membranes were blocked in 5% non-fat milk in TBST, followed by immunoblotting with the following antibodies: PHGDH (Sigma Aldrich, HPA021241-100), GFP (Cell Signaling, 2956), HSP90 (Cell Signaling, 4874), β-ACTIN (Thermo Fisher, AM4302, clone AC-15).

### Statistical analysis

Data were analyzed using a two-sided unpaired Student’s t test or Kaplan-Meier analysis as noted. GraphPad Prism 7 and 8 software were used for all statistical analyses, and values of p<0.05 were considered statistically significant (*p<0.05; **p<0.01; ***p<0.001).

## Results

### Generation of an inducible model for systemic PHGDH knockdown

In order to evaluate the requirement for serine biosynthesis in normal, proliferating adult tissues, we generated a mouse model in which an shRNA targeting PHGDH is linked to GFP and expressed in a doxycycline-inducible manner (**Figure 1A**). To generate this model, we first tested the efficacy of single copy shRNAs targeting mouse PHGDH in NIH3T3 cells (**Supplementary Figure 1**). shRNA #3 was selected, cloned into a col1A1 targeting vector for recombination-mediated cassette exchange in embryonic stem cells, and used to generate shPHGDH mice. Mice expressing Renilla Luciferase targeting shRNA (shREN) were used as controls (Premsrirut et al., 2011).

**Figure 1:**
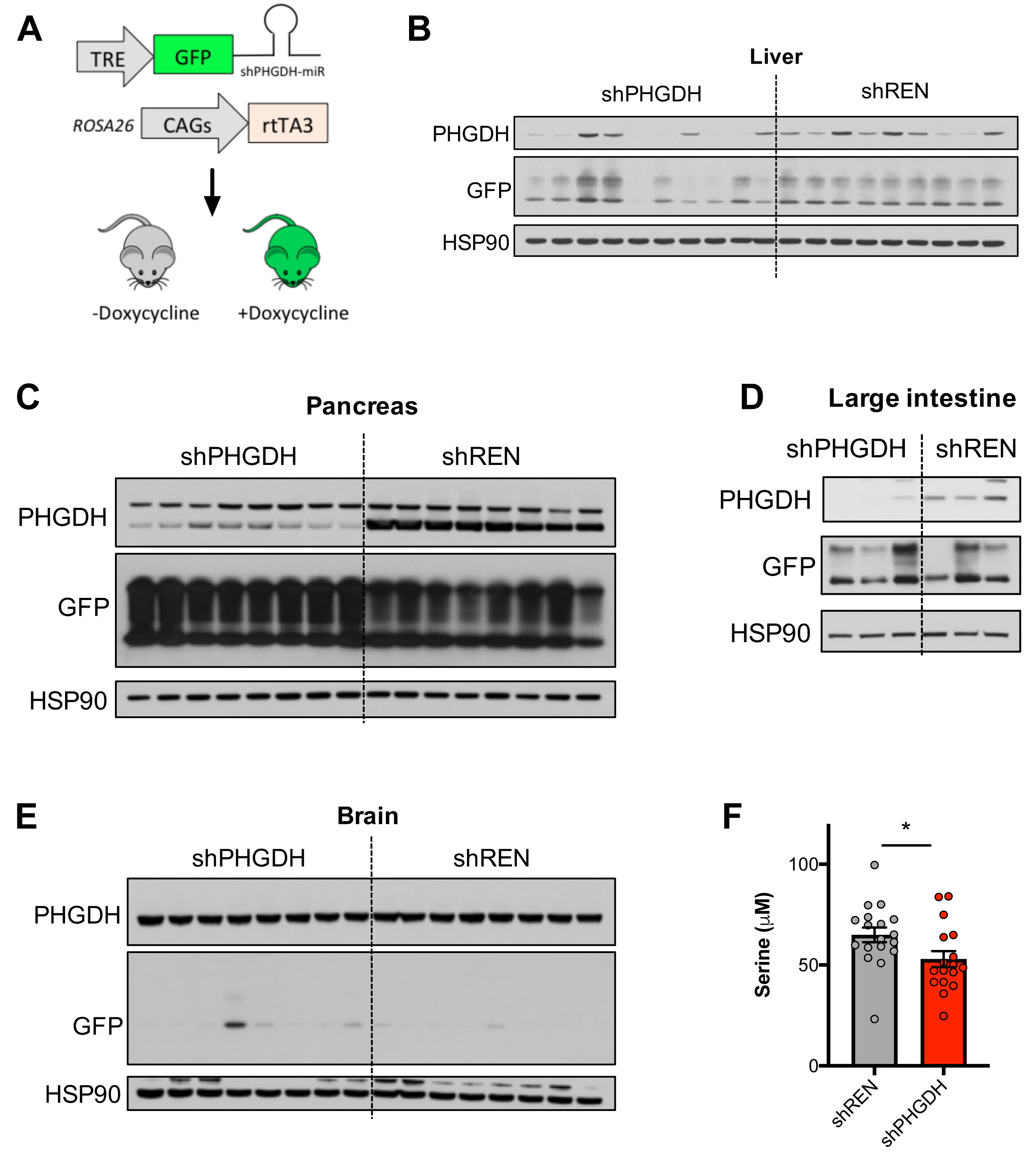
**Generation of an inducible model for systemic PHGDH knockdown** (A) Schematic representation of the inducible shRNA system. (B-E) Western blot analysis of PHGDH, GFP and HSP90 protein levels in the liver (B), pancreas (C), large intestine (D) and brain (E) of shPHGDH and shREN mice. Mice were placed on a 200ppm doxycycline diet for 4-8 months. (F) Serum serine concentrations of 8-month-old shREN (N=16) and shPHGDH (N=16) mice.

shREN and shPHGDH mice were crossed with a ubiquitous reverse tetracycline transactivator allele (rtTA3) to allow for whole body expression of the shRNA upon doxycycline administration. We selected the CAGs-rtTA3 allele, which demonstrated robust rtTA3 expression and activity in the pancreas, liver, kidney, small intestine, large intestine, skin, thymus and bone marrow, although limited activity was observed in the spleen (Dow et al., 2014). Further, because doxycycline levels achieved in mouse brain are an order of magnitude lower than in plasma (Lucchetti et al., 2019), this model is expected to spare the brain and avoid the previously reported brain toxicity associated with PHGDH knockout. shRNA; rtTA3 dual allele mice were placed on a 200 ppm doxycycline-containing diet containing serine and glycine. As expected, western blot analysis of PHGDH expression in tissues revealed that PHGDH knockdown was achieved in the pancreas, liver, and large intestine (Figures 1B-D), but was absent in the brain and spleen (**Figure 1E**, **Supplemental Figure 2**). Interestingly, we observed significant variation in liver PHGDH between animals, which may be due to circadian influences as tissues were not collected at a specific time of day. Serum serine levels were decreased by approximately 20% following PHGDH knockdown (**Figure 1F**), with the remaining 80% likely accounted for by dietary serine and glycine. These results demonstrate that our inducible PHGDH knockdown model spares brain and spleen PHGDH but diminishes PHGDH in other tissues where rtTA3 is robustly expressed, resulting in a reduction in circulating serine.

### PHGDH is not required for tissue proliferation and mouse viability

We next assessed the consequence of PHGDH depletion on animal health and viability. Mice expressing shPHGDH were found to have a normal lifespan, with only a modest, non-significant reduction in survival compared to shREN mice (**Figure 2A**). Because PHGDH has been associated with proliferation in neoplastic cells, we determined the effects of PHGDH silencing on the most proliferative tissues in adult mice. First, we examined the health of the intestine, which contains highly proliferative stem cells in the crypt, which must replace the entire epithelium every few days. We found that mice gained weight at normal rates, and intestines exhibited a normal morphology and proliferation rate (Figure 2B,C). Next, we examined the hematopoietic cells in the bone marrow. We found no overtly abnormal phenotypes in the bone marrow of shPHGDH mice in the absence of stress (**Figure 2D**). Further, shPHGDH mice had normal red blood cell and white blood cell counts (Figure 2E,F). However, this may be explained by low basal PHGDH expression in the bulk bone marrow population, which was undetectable by western blot (not shown). Finally, because PHGDH knockdown was very robust in the pancreas, we examined the consequence of PHGDH depletion on glucose tolerance. shREN and shPHGDH mice were administered a bolus of glucose in an oral glucose tolerance test and blood glucose was assayed over time. We found that glucose tolerance was not affected by PHGDH knockdown (**Figure 2G**), suggesting that PHGDH is not required for normal pancreatic function. Further, PHGDH knockdown in the liver did not affect liver function as determined by blood markers for liver enzymes and other liver markers (**Supplementary Figure 3**). Collectively, these results demonstrate that PHGDH is not required for the cellular proliferation or normal function of multiple tissues in adult mice, which present with no overtly abnormal phenotypes upon PHGDH knockdown.

**Figure 2:**
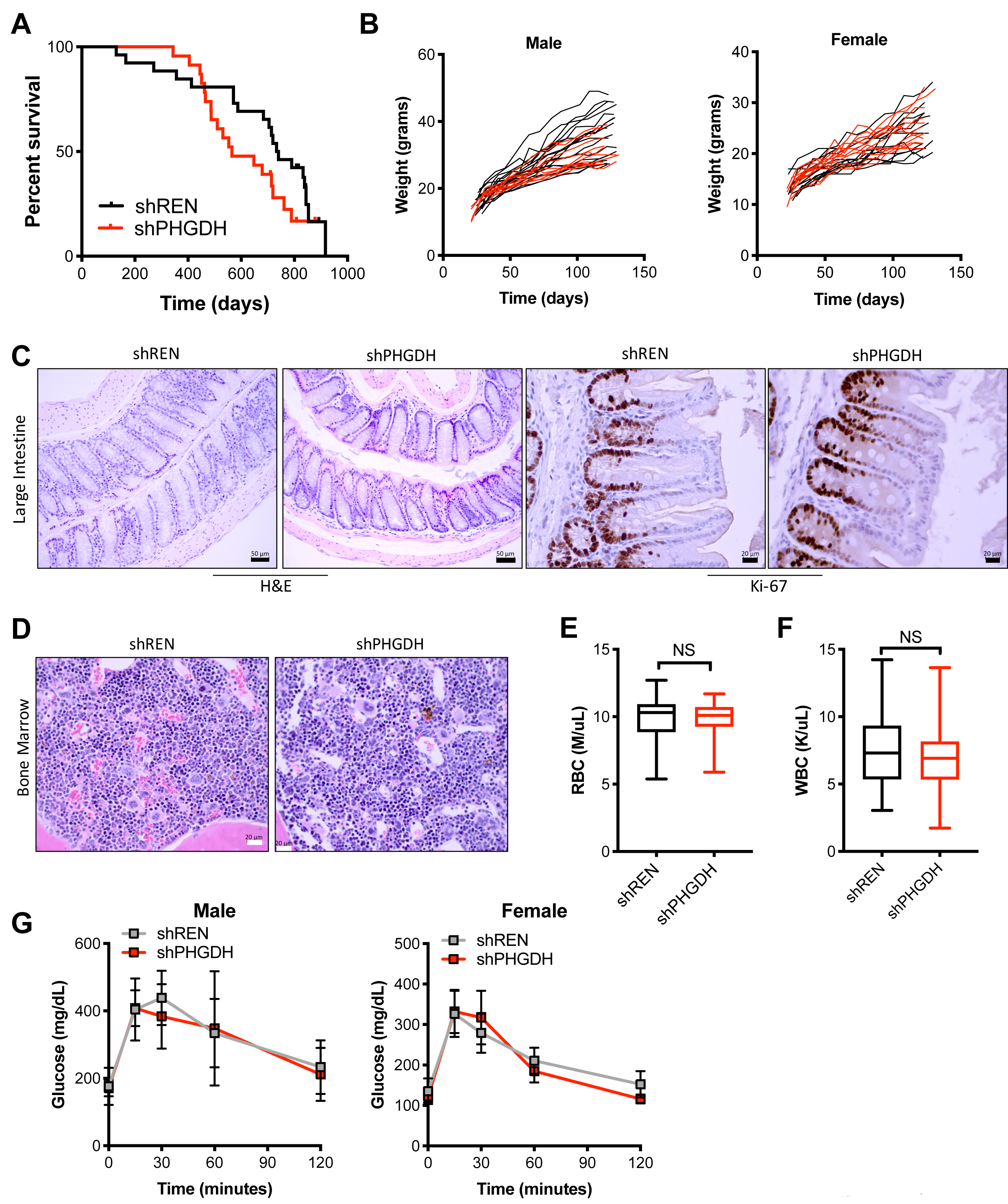
**Systemic PHGDH depletion is non-toxic.** (A) Overall srvival of mice expressing shPHGDH (N=23) or the control shRenilla (shREN, N=26). (B) Weight of male and female mice expressing shPHGDH (male, N=16; female, N=15) or shREN (male, N=13; female, N=10). Mice were placed on doxycycline at weaning. (C) Representative hematoxylin & eosin stained (N=10+) and Ki-67 immunostained (N=5 each) large intestine sections from shPHGDH and shREN mice at endpoint. (D) Representative hematoxylin & eosin stained bone marrow sections from shPHGDH and shREN mice (N=10+) at endpoint. (E) Red blood cell counts of shREN (N=31) and shPHGDH (N=33) mice at 8 months. (F) White blood cell counts of shREN (N=31) and shPHGDH (N=33) mice at 8 months. (G) Oral glucose tolerance test at 100 days on doxycycline. Male and female shPHGDH and shREN mice were challenged with 2g/kg glucose at time=0 and blood glucose levels were assayed at the indicated time points. Male shPHGDH (N=7), Male shREN (N=7), Female shPHGDH (N=6), Female shREN (N=10).

### shPHGDH mice do not exhibit deoxysphingolipid formation

In addition to the importance of serine for the proliferation of neoplastic and non-neoplastic cells, low serine levels have been linked with toxic deoxysphingolipid formation in both serine deprivation experiments (Gantner et al., 2019) and PHGDH knockout brains (Esaki et al., 2015). When serine levels are low, deoxysphingolipids are made via serine palmitoyltransferase (SPT) using alanine as a substrate instead of serine, thereby leading to deoxysphingolipid accumulation and cellular toxicity, particularly in the central nervous system. To examine the effect of PHGDH depletion on deoxysphingolipid metabolism, we first examined deoxysphingolipid levels in the circulation. However, we found no accumulation of individual (**Figure 3A**) or total (**Figure 3B**) dihydro-deoxy (dihydro-dox) or the deoxy (dox) sphingolipid species in the serum of shPHGDH mice. Similarly, liver deoxysphingolipid levels were not altered (**Figure 3C**), with the levels of dihydro-deoxysphingolipids actually significantly lower upon PHGDH knockdown (**Figure 3D**). Finally, the levels of the deoxysphingolipid precursor deoxysphinganine was not elevated in liver (**Figure 3E**). These results demonstrate that the decrease in circulating serine following PHGDH knockdown is not sufficient to induce deoxysphingolipid formation.

**Figure 3:**
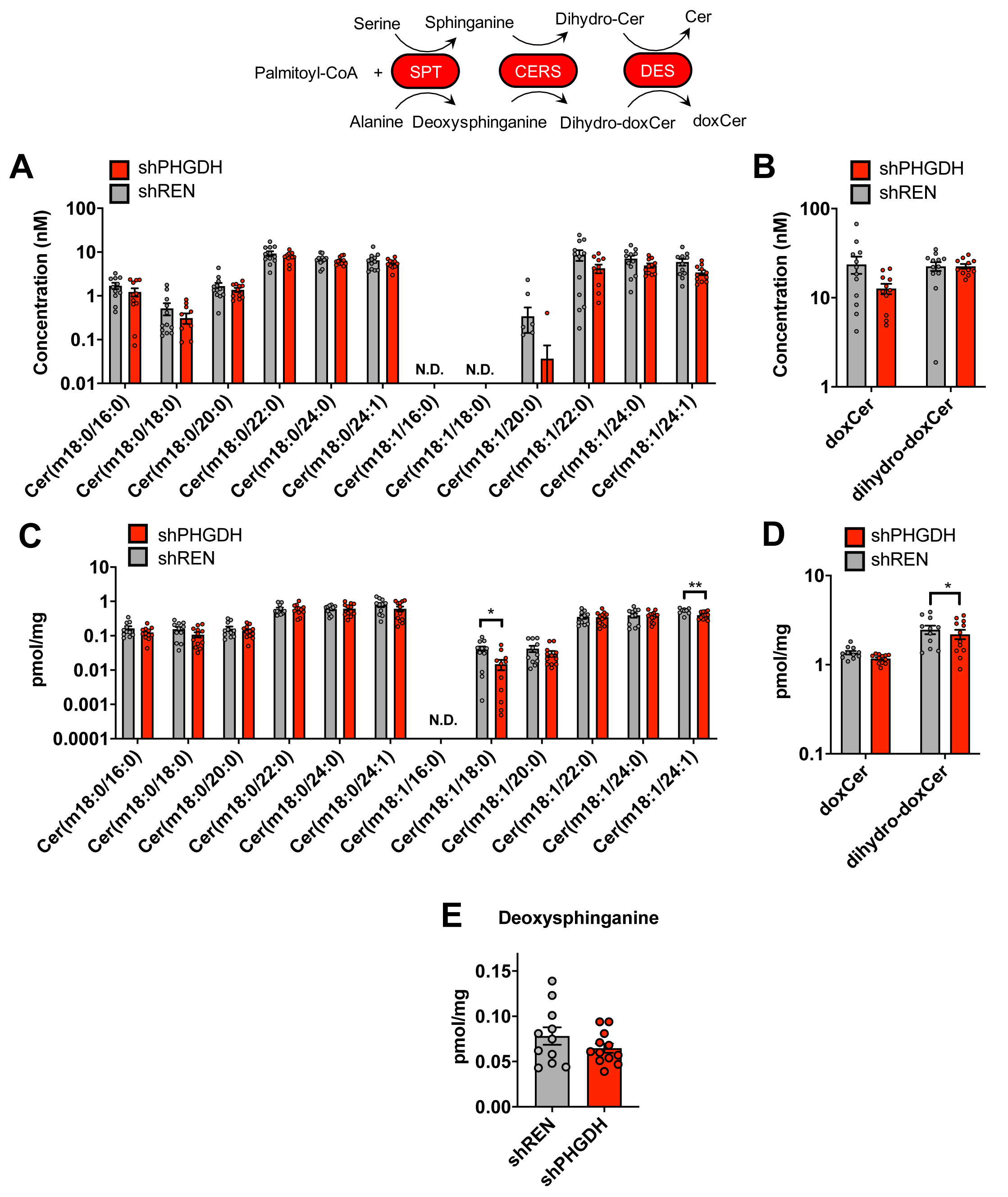
**PHGDH knockdown does not promote deoxyshingolipid formation but depletes ceramides** (A) Concentration of individual deoxysphingolipids and dihydrodeoxysphingolipids in the serum of shREN and shPHGDH mice. (B) Total concentration of deoxysphingolipids (doxCer) and dihydrodeoxysphingolipids (dihydro-doxCer) in the serum of shREN and shPHGDH mice. (C) Quantity of deoxysphingolipids in the liver of shREN and shPHGDH mice. Quantities were normalized to mg of tissue. (D) Total quantity of deoxysphingolipids (doxCer) and dihydrodeoxysphingolipids (dihydro-doxCer) in the liver of shREN and shPHGDH mice. Quantities were normalized to mg of tissue. (E) Quantity of deoxysphinganine in the liver of shREN and shPHGDH mice. Quantities were normalized to mg of tissue. For A-E: 8-month-old shREN (N=11) and shPHGDH (N=12) mice were used for analysis.

### PHGDH knockdown decreases serum and liver ceramides

Importantly, the canonical product of the SPT pathway, ceramide, can also be influenced by serine availability (Gao et al., 2018). Interestingly, and in agreement with a previous report demonstrating that serine limitation depletes ceramide (Gao et al., 2018), we observe a decrease in many ceramide species in the serum of shPHGDH mice (**Figure 4A**). Consistent with what was observed in the serum, liver ceramides were also significantly decreased (**Figure 4B**). Analysis of liver serine levels revealed a decrease by approximately 20% (**Figure 4C**), similar to what was observed in the serum (**Figure 1E**). Collectively, these results demonstrate that PHGDH knockdown influences ceramide levels in vivo, consistent with prior studies describing effects of serine availability on ceramide metabolism.

**Figure 4:**
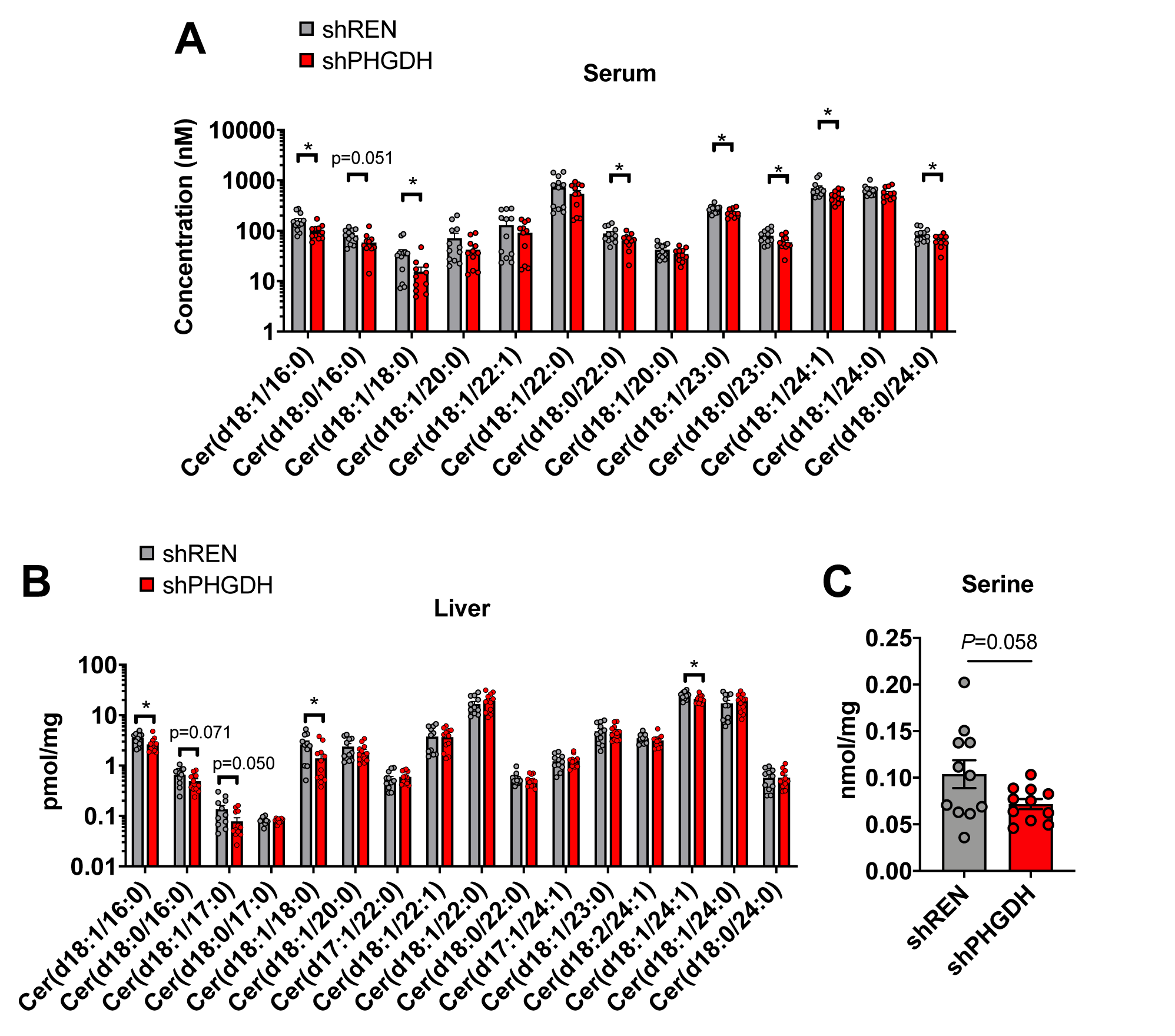
**PHGDH knockdown depletes ceramides** (A) Concentration of individual ceramides in the serum of 8-month-old shREN (N=12) and shPHGDH mice (N=11). (B) Quantity of individual ceramides in the liver of 8-month-old shREN (N=11) and shPHGDH (N=12) mice. Quantities were normalized to mg of tissue. (C) Liver serine quantities of 8-month-old shREN (N=11) and shPHGDH (N=11) mice. Quantities were normalized to mg of tissue.

**Figure 5:**
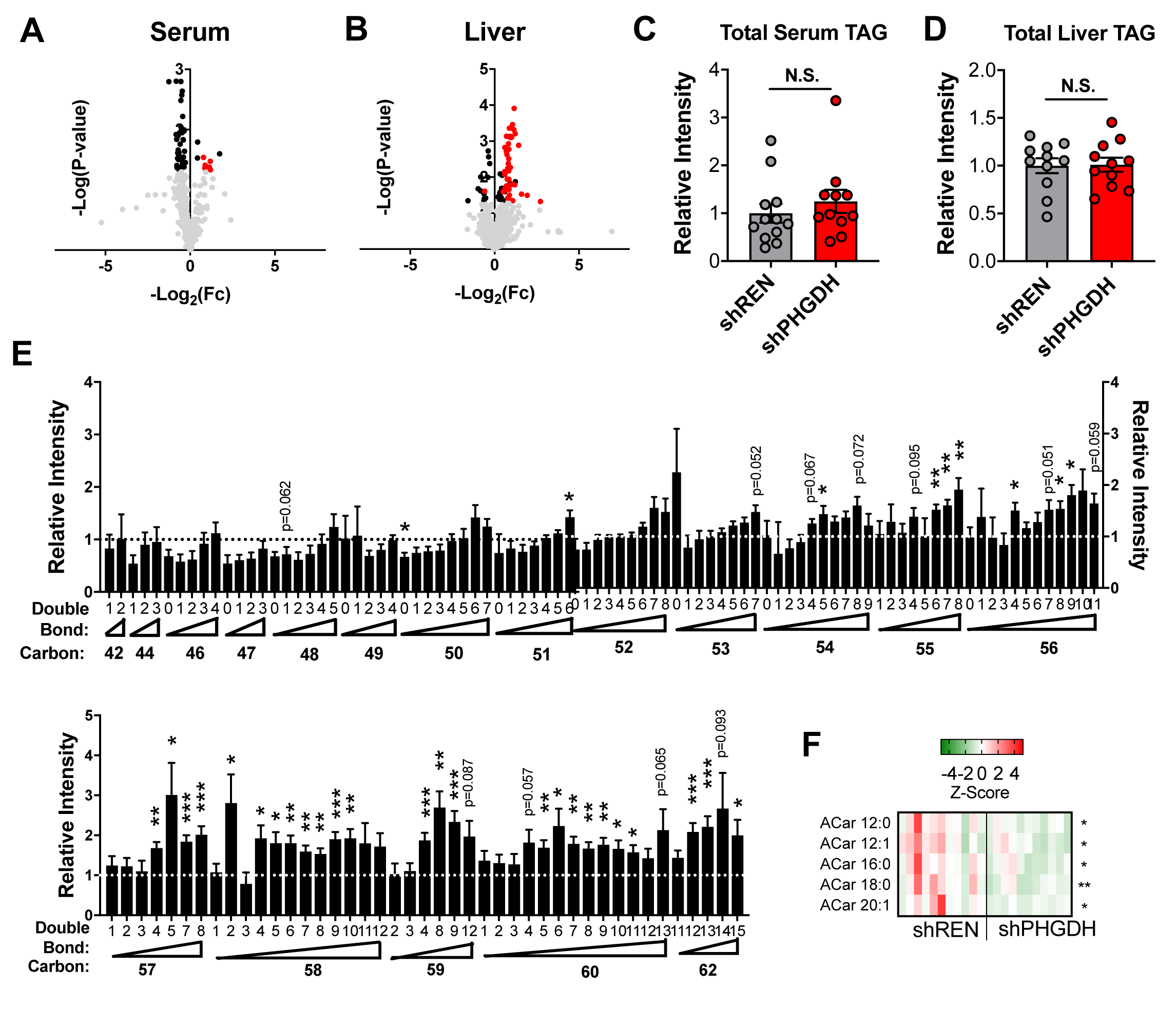
**PHGDH knockdown alters liver lipid profiles** (A) Volcano plot of lipidomics analysis of shPHGDH (N=11) serum compared to shREN (N=12). Significant metabolites are in bold. Triacylglycerol species are indicated in red. (B) Volcano plot of lipidomics analysis of shPHGDH (N=11) liver compared to shREN (N=11). Significant metabolites are in bold. Triacylglycerol species are indicated in red. (C-D) Total triacylglycerol (TAG) levels in the serum (C) and liver (D) of shREN and shPHGDH mice. Levels are normalized to shREN. (E) Individual TAG species in the liver of shPHGDH mice compared to shREN. Levels are normalized to shREN. (F) Fatty acyl carnitine (Car) levels in the liver of shREN and shPHGDH mice from the analysis in (B). Only significant (p < 0.05) metabolites are shown.

### PHGDH knockdown alters triacylglycerol composition

We next examined whether impaired sphingolipid synthesis influenced other lipid classes. Multiple lipid classes, such as sphingolipids, glycerophospholipids, and triacylglycerols, share similar fatty acid tails, with distinct lipid head groups. Lipidomic analysis of shREN and shPHGDH serum and liver revealed global lipid alterations (Figure 4A-B, Supplementary **Tables 2** and **3**), suggesting a perturbation in lipid metabolism following PHGDH knockdown. In particular, we observed a significant accumulation of certain triacylglycerol (TAG) species, particularly in the liver (**Figure 4B**). However, total TAG levels in the serum and liver were unchanged (Figure 4C-D). Rather, it was the composition of the TAG species that was altered. Specifically, accumulated TAG species were enriched in long chain (LC) polyunsaturated fatty acid (PUFA) tails, while short chain saturated TAGs were decreased (**Figure 4E**). Further, lipidomics analysis of fatty acyl carnitine species revealed a significant decrease in C12:0-, C12:1-, C16:0-, C18:0-, and C20:1-carnitine species, suggesting a decrease in fatty acids availability for β-oxidation (**Figure 4F**). These results demonstrate that PHGDH knockdown alters fatty acyl-carnitine levels and TAG composition but does not affect total TAG levels.

## Discussion

Genetic deficiency in the de novo serine biosynthesis genes *PHGDH*, *PSAT1*, and *PSPH,* in humans causes Neu-Laxova syndrome (NLS), a very rare autosomal recessive congenital disorder (Acuna-Hidalgo et al., 2014; Shaheen et al., 2014). Severity is dictated by the degree of pathway activity loss and most patients are affected from infancy. NLS patients present with central nervous system (CNS) symptoms including microcephaly, impaired motor function, epilepsy, and perinatal lethality. By contrast, we find that PHGDH depletion in non-cerebral tissues following the post-natal period results in no overt phenotype, which is consistent with the symptoms of PHGDH deficiency being predominantly localized to the CNS.

Our model is useful for the study of PHGDH in adult tissues, but knockdown is not complete in all tissues studied. While we could achieve better knockdown in some tissues with a diet containing 625ppm doxycycline (not shown), concerns about the effects of doxycycline on metabolism led us to choose a lower concentration. In addition, improved PHGDH depletion in some tissues can likely be achieved through the use of other rtTA3 alleles that have higher promoter activity in those tissues. Consequently, caution must be used when interpreting our results on the effect of PHGDH knockdown in all tissues because it is possible some cell types still have abundant PHGDH expression due to poor shRNA expression. Despite this limitation, our model is likely to more accurately model the effects of PHGDH inhibition in select tissues compared to whole body knockout due to the incomplete and transient nature of inhibition of enzymes by small molecules.

Our findings suggest that PHGDH inhibitors that cannot cross the blood brain barrier may be well tolerated, provided adequate serine and glycine are supplied through the diet. While liver lipid metabolism was altered, this did not appear to induce any pathology in the absence of stress. Our findings demonstrating that PHGDH knockdown results in a depletion of ceramides are very similar to findings seen with serine starvation in colon cancer cells, where ceramide depletion also led to loss of mitochondrial function, suggesting that mitochondrial function may also be impaired in our model as well. Interestingly, inhibition of mitochondrial function was recently found to induce the synthesis of highly unsaturated fatty acids (HUFA) to recycle glycolytic NAD+ (Kim et al., 2019). Alternatively, altered TAG metabolism in our model may be a consequence of decreased Palmitoyl-CoA utilization for sphingolipid synthesis. Fatty acid incorporation into TAGs has been shown to protect against lipotoxicity in many settings (Brookheart et al., 2009; Listenberger et al., 2003). Further work is needed to determine whether shPHGDH livers have mitochondrial impairment, and whether alterations in TAG metabolism protect shPHGDH livers.

## Conclusions

Our study found that PHGDH knockdown has a modest effect on circulating serine in the presence of dietary serine/glycine and does not affect the function or proliferation of adult pancreas, liver, and intestine. However, loss of PHGDH expression in the liver reduced ceramide levels without increasing the levels of deoxysphingolipids. Further, liver triacylglycerol profiles were altered, with an accumulation of longer-chained, polyunsaturated tails upon PHGDH knockdown. Collectively, these results suggest that PHGDH-derived serine supports liver ceramide synthesis and sustains general lipid homeostasis.

## List of abbreviations

CNS: central nervous system
dihydrodox: dihydrodeoxy
dox: deoxy
NAD+: nicotinamide adenine dinucleotide
NLS: Neu-Laxova syndrome
PHGDH: D-3-phosphoglycerate dehydrogenase
shPHGDH: shRNA targeting PHGDH
shREN: shRNA targeting Renilla luciferase
shRNA: small hairpin RNA
SPT: serine palmitoyltransferase

## Declarations

### Ethics approval and consent to participate

Mice were housed and bred in accordance with the ethical regulations and approval of the IACUC (protocol #IS00003893R).

### Consent for publication

Not applicable.

### Availability of data and materials

All data generated or analyzed during this study are included in this published article and its supplementary information files. Materials are available from the corresponding author on request.

## Competing interests

The authors declare that they have no competing interests.

## Funding

GMD is supported by grants from the NIH (R37-CA230042), and the PanCAN/AACR Pathway to Leadership Award. YPK is supported by AACR-Takeda Oncology Lung Cancer Research Fellowship. FAK is supported by grants from the NIH (K22-CA197058 and R03-CA227349). The Proteomics/Metabolomics Core is supported in part by the NCI (P30-CA076292), Moffitt Foundation, and a Florida Bankhead-Coley grant (06BS-02-9614). These funding bodies had no role in the design of the study and collection, analysis, and interpretation of data, or in writing the manuscript.

## Author contributions

GMD designed the study, FAK and GMD generated the shPHGDH mouse, AF performed animal experiments and analyses, JJS performed pathology analyses, YPK and ML established lipidomics methodology, YPK performed metabolomics, lipidomics and western blotting, GMD wrote the manuscript and all authors commented on it.

## Acknowledgements

We thank Lukas Dow for providing the shREN and CAGs-rtTA3 mice and for advice on the cloning, ESC targeting, and generation of shPHGDH mice.

---

**Supplementary Figure 1.**
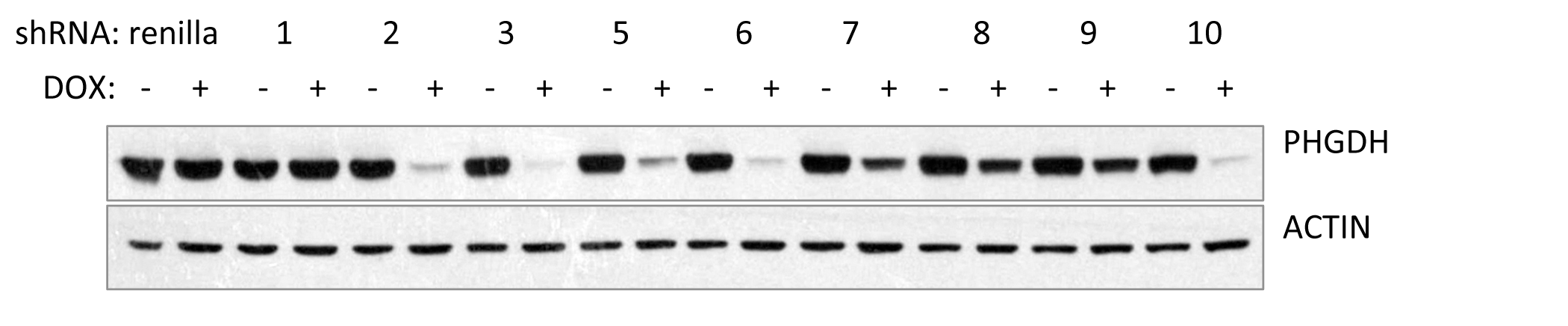
**Validation of shRNAs for model development** NIH3T3 cells expressing Renilla or PHGDH-targeting shRNAs (#1-10) were treated with 1μg/mL doxycycline for 6 days and PHGDH expression determined by western blot. β-actin is used as a loading control.

**Supplementary Figure 2.**
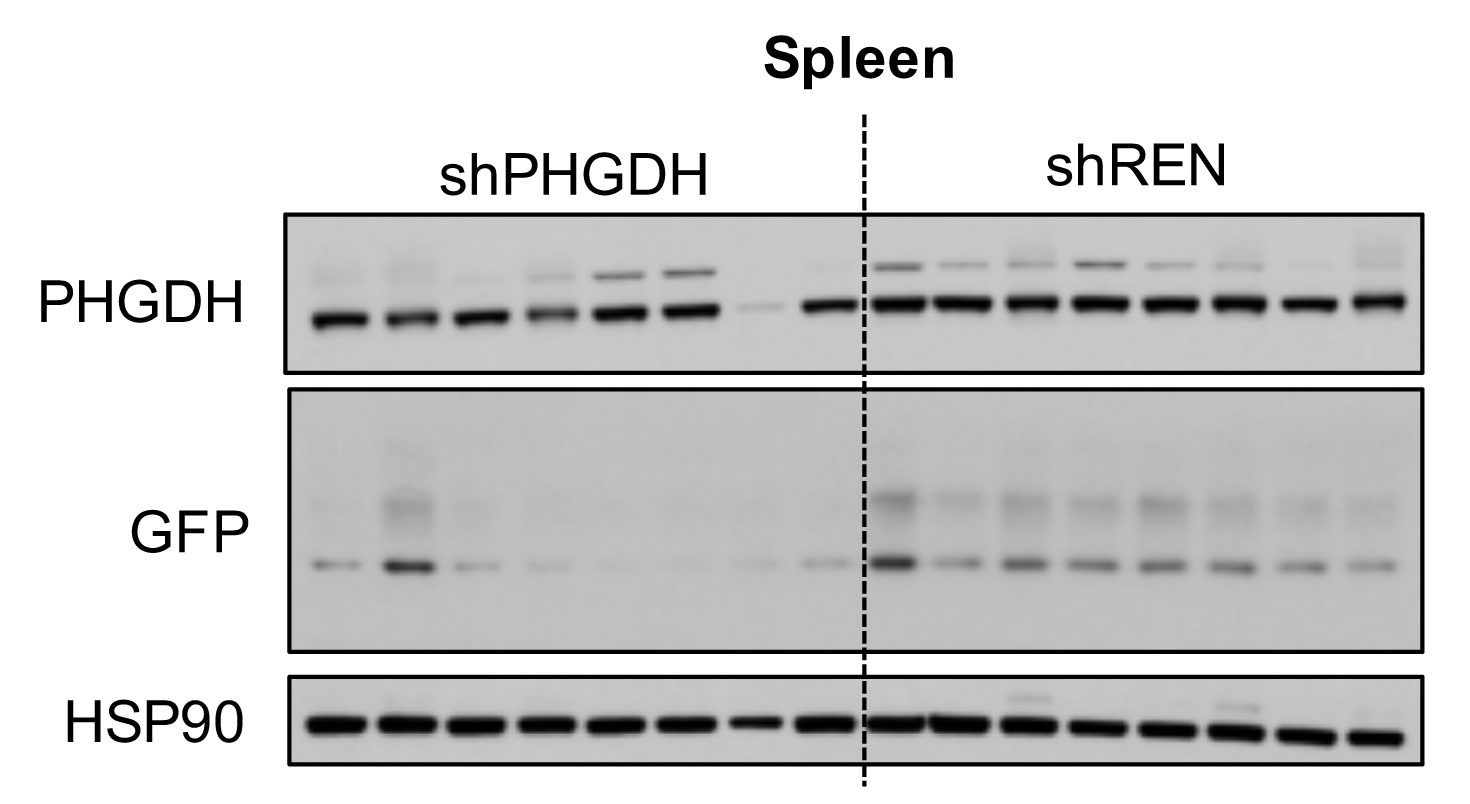
**shPHGDH have poor knockdown in the spleen** Western blot analysis of PHGDH, GFP and HSP90 protein levels in spleen of shPHGDH and shREN mice. Mice were placed on a 200ppm doxycycline diet for 8 months.

**Supplementary Figure 3.**
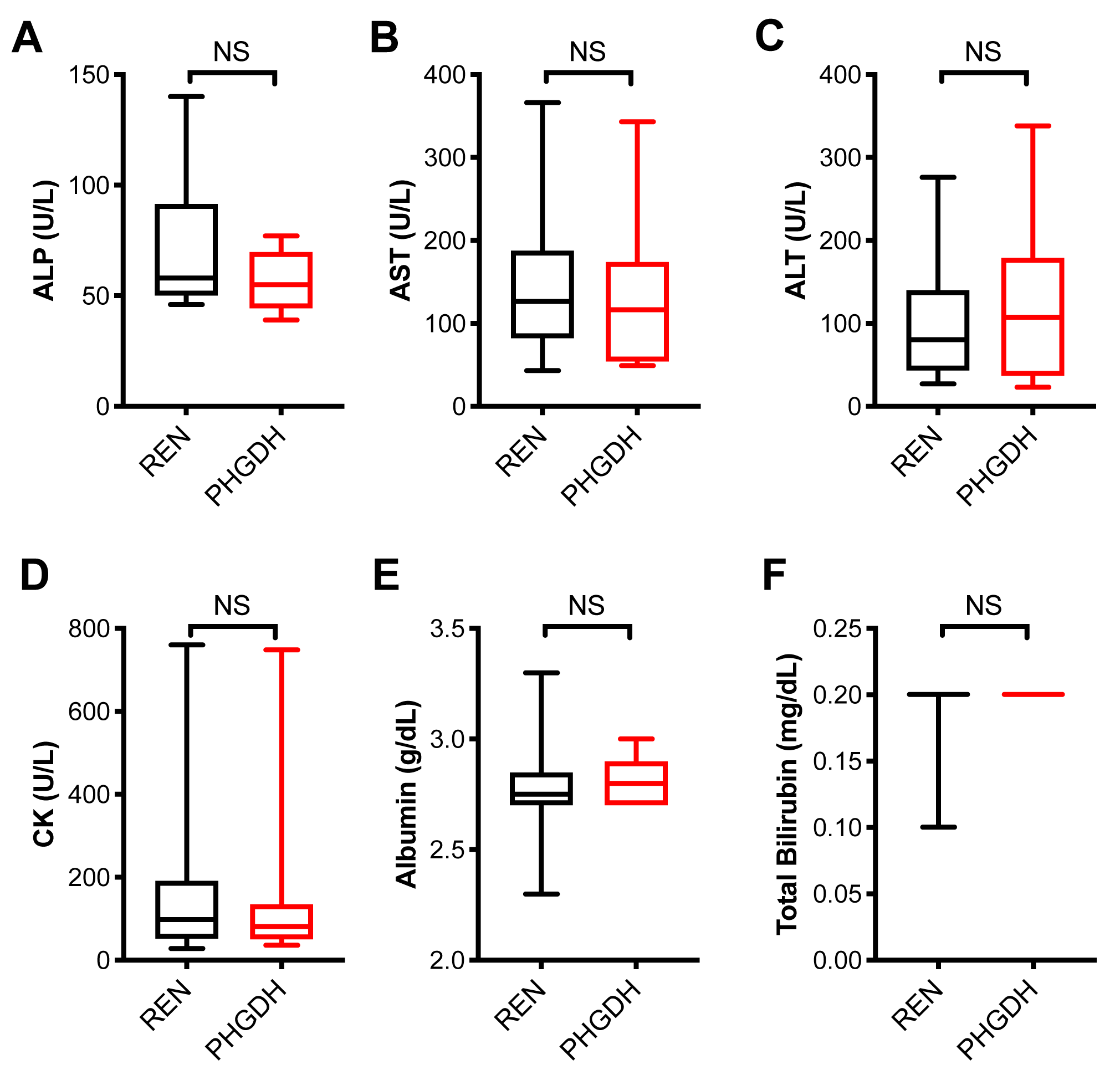
**PHGDH knockdown does not affect liver function** shPHGDH (N=10) and shREN (N=10) mouse serum was collected and analyzed by IDEXX (Liver Panel). (A) ALP – Alkaline phosphatase. (B) AST – aspartate transaminase. (C) ALT – alanine transaminase. (D) CK – creatine kinase. (E) – Total albumin. (E) Total bilirubin.

**Supplementary Table 1.**
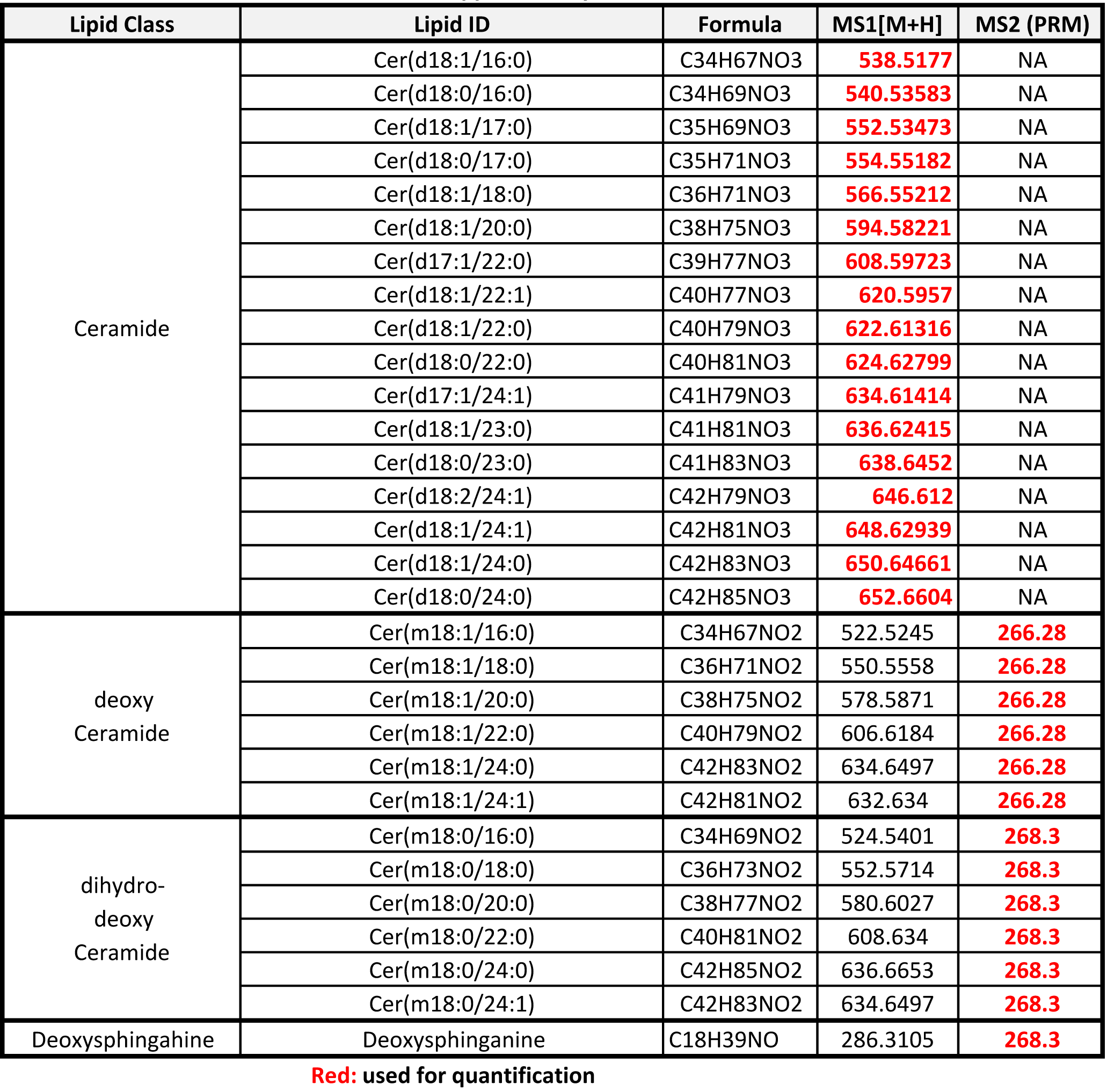
Analysis parameters for targeted lipidomics

**Supplementary Table 2.**
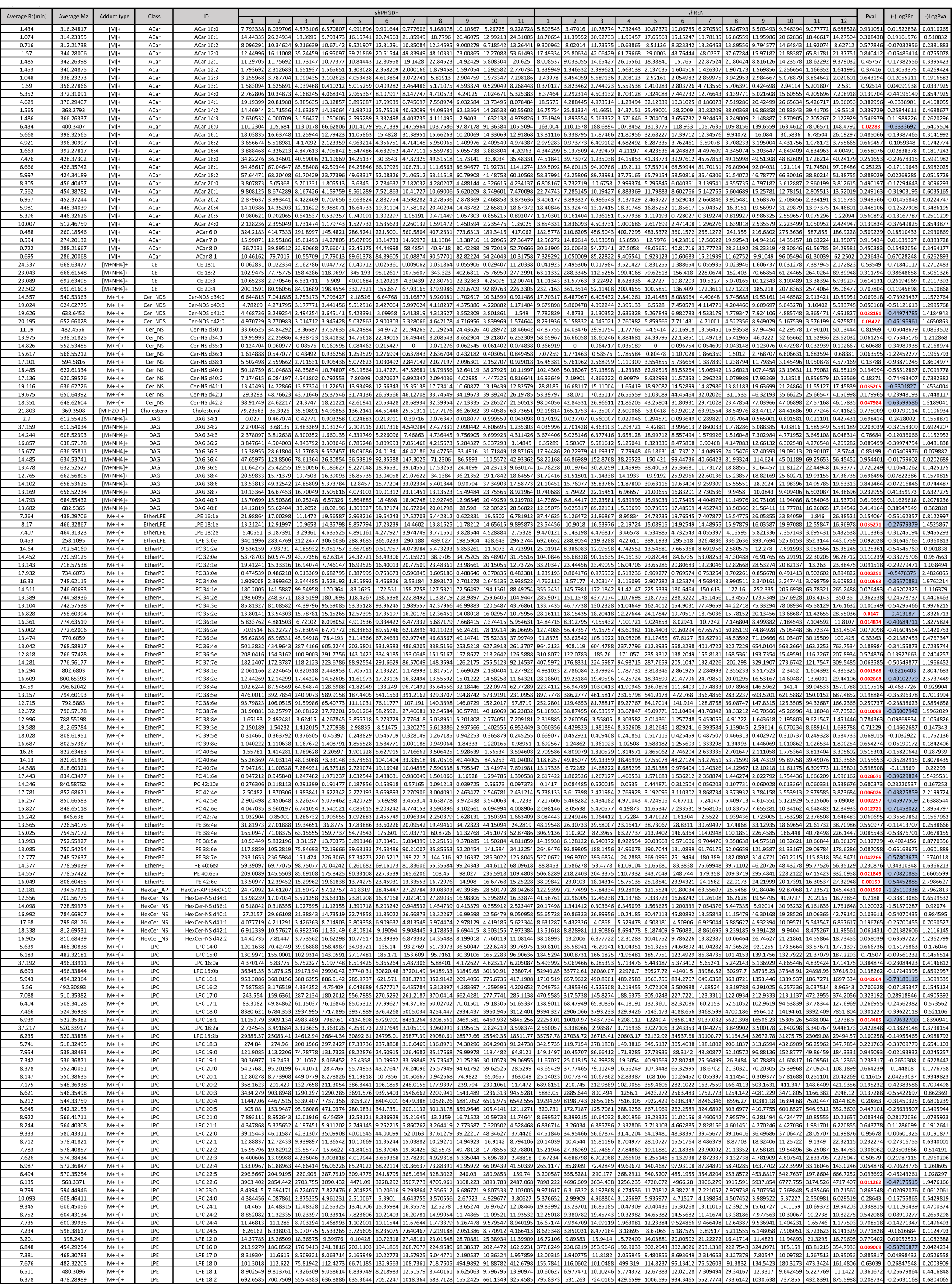
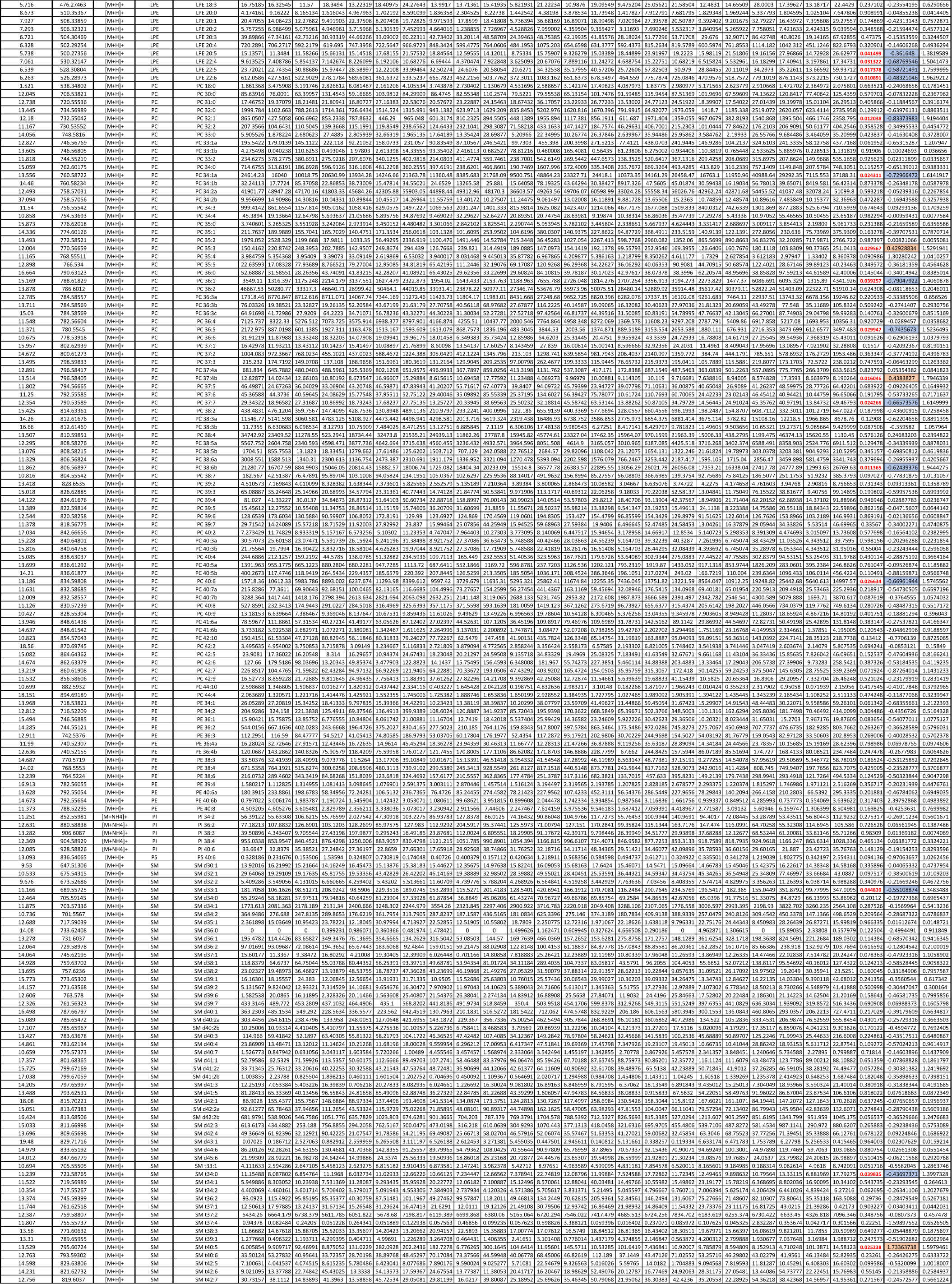
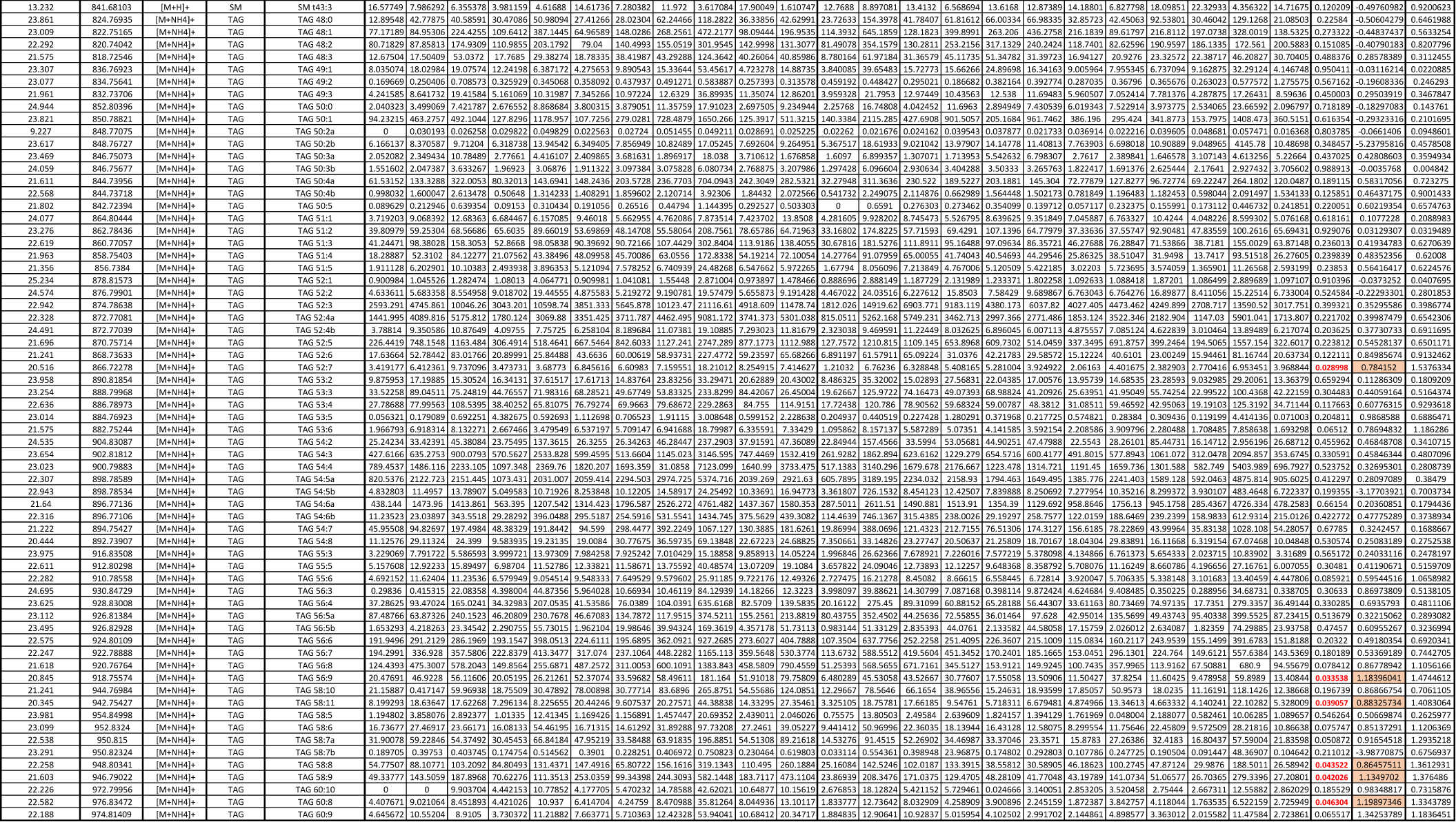
Lipidomics data from Figure 5A. Lipidomics analysis of shPHGDH (N=11) serum compared to shREN (N=12).

**Supplementary Table 3.**
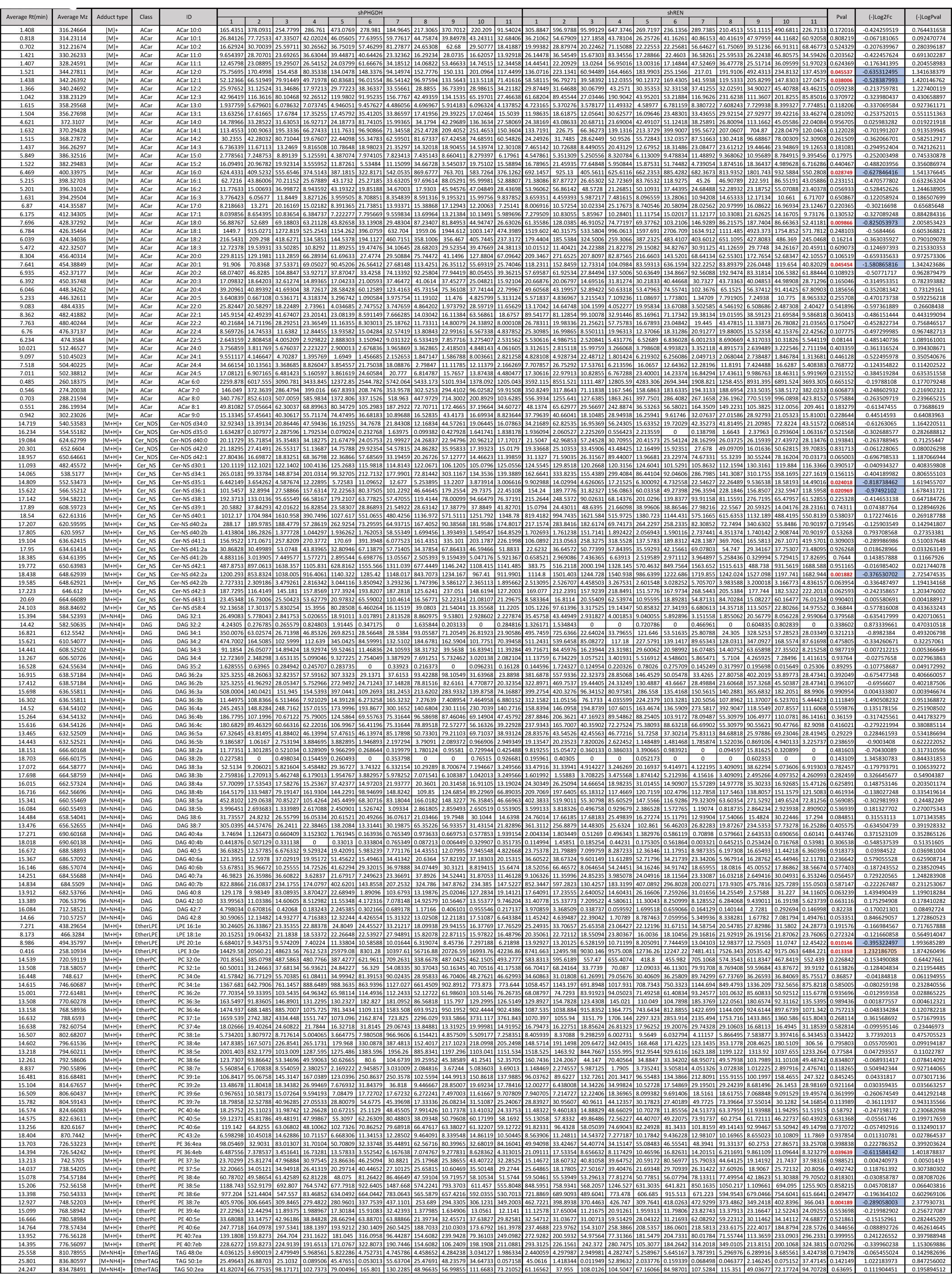
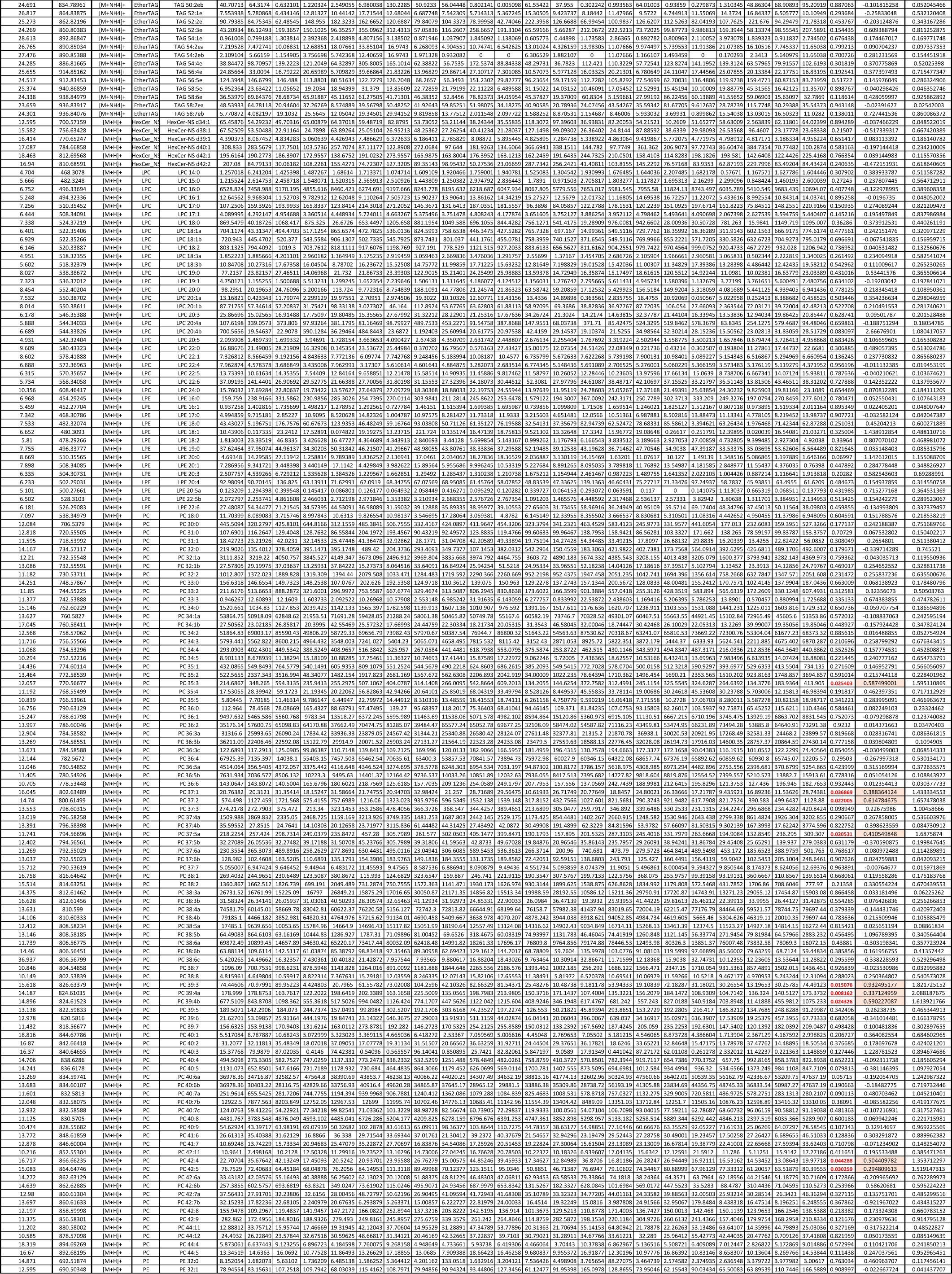
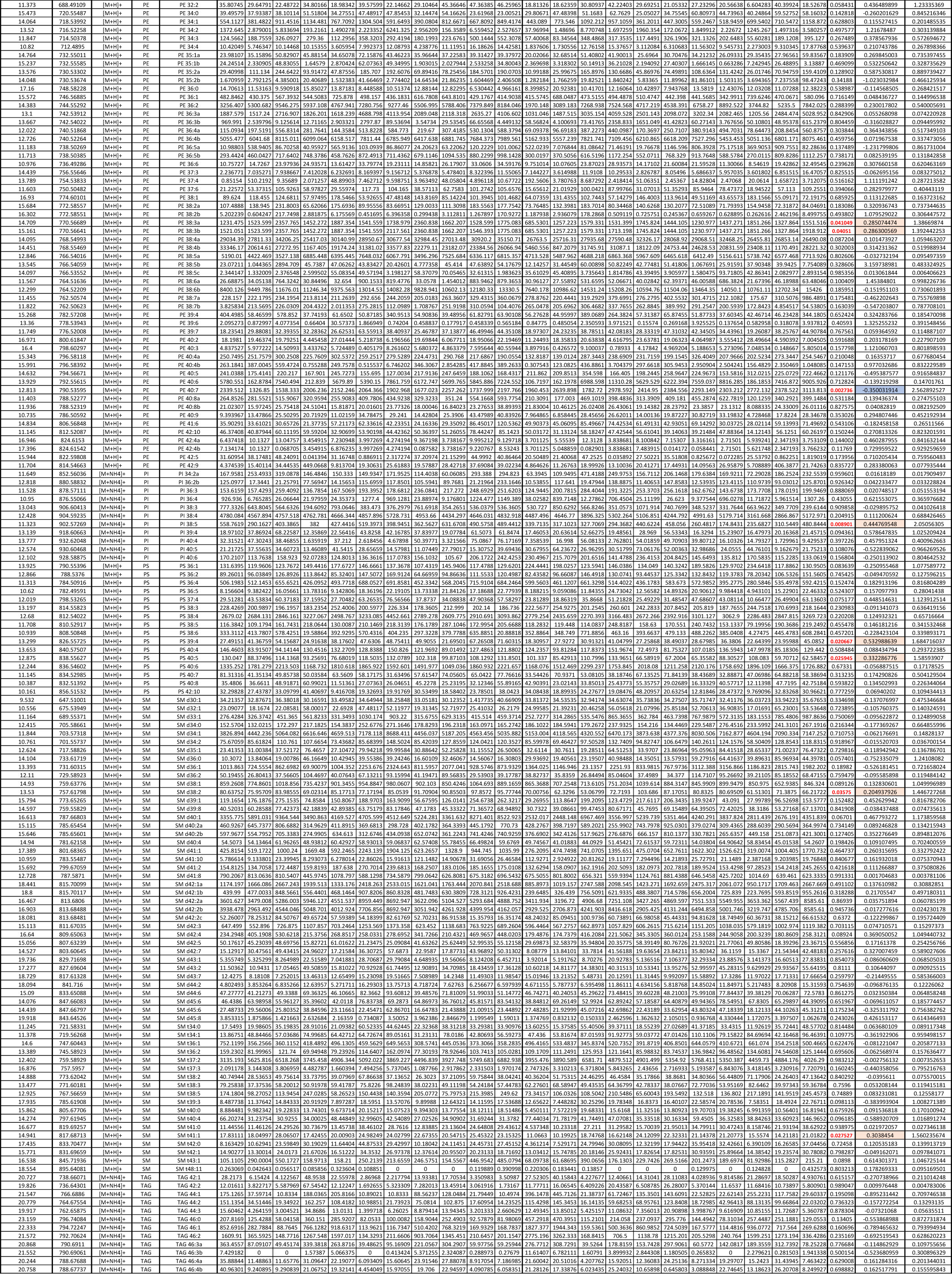
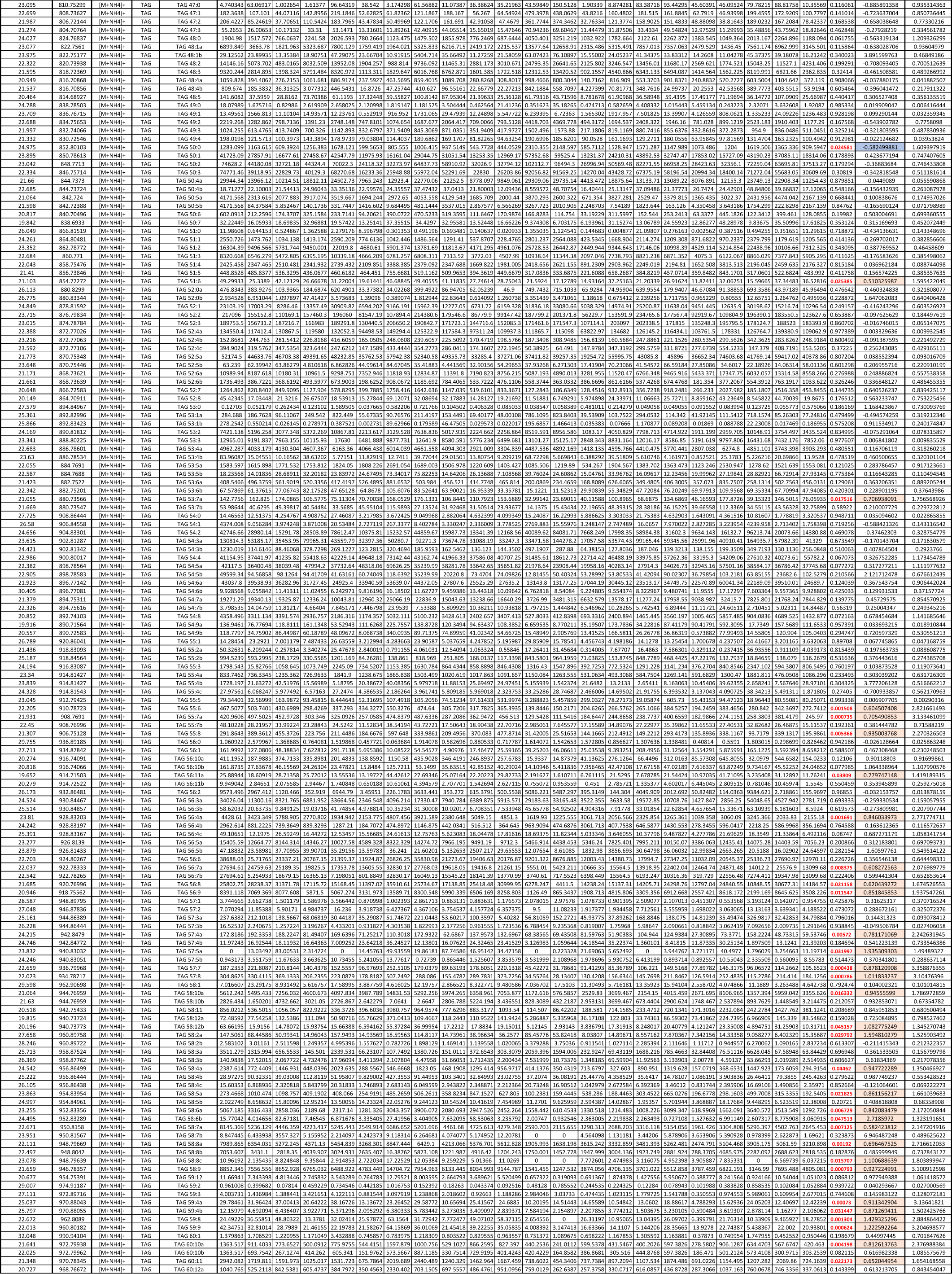
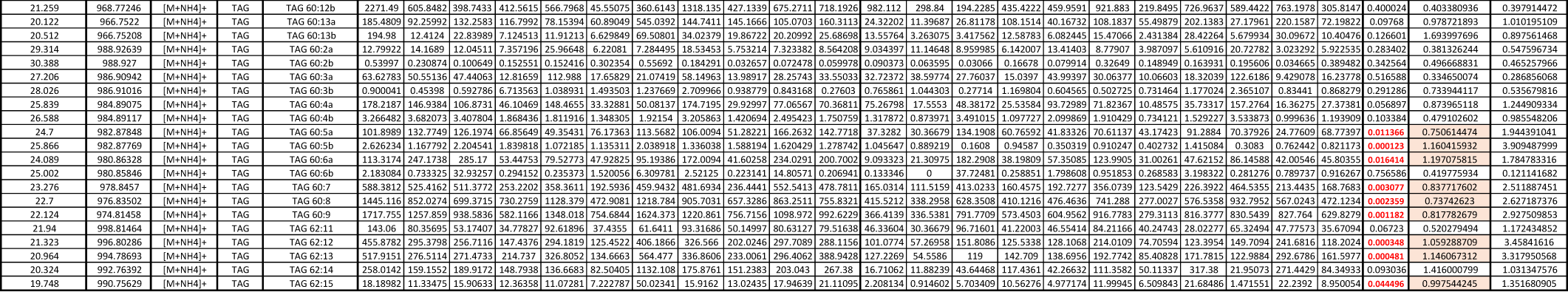
Lipidomics data from Figure 5B. Lipidomics analysis of shPHGDH (N=11) liver compared to shREN (N=11).

